# The effect of task demand on EEG responses to irrelevant sound and speech in simulated surgical environments

**DOI:** 10.1101/2025.02.13.638036

**Authors:** Marc Rosenkranz, Verena N. Uslar, Dirk Weyhe, Martin G. Bleichner

## Abstract

Complex soundscapes in high-stakes environments, such as the operating room (OR), are characterized by a variety of overlapping auditory stimuli and present significant challenges for personnel, particularly during periods of high demand. This study investigates how task demand and an OR soundscape including irrelevant speech, influence perceived workload, surgical performance, and auditory processing in a simulated surgical environment, using mobile electroencephalography (EEG). Participants performed two simulated surgical tasks, namely peg transfer and suturing, representing a low-demand and high-demand task, respectively. The tasks were performed under two sound conditions: An OR soundscape was presented with irrelevant speech or alone. Neural responses to transient and continuous auditory stimuli were analyzed using event-related potentials (ERPs) and temporal response functions (TRFs), respectively. Results showed that irrelevant speech increased self-reported workload and distraction. EEG analyses revealed reduced neural responses to transient sounds and irrelevant speech under high task demand, reflecting early-stage sensory filtering of auditory distractions. Notably, an inverse relationship was observed between neural responses to speech and self-reported workload, indicating that the speech responses may serve as a marker for perceived workload. Overall, this study demonstrates the potential of EEG to assess irrelevant sound processing in realistic work-like settings and highlights the critical role of task demand in modulating neural responses and self-reported workload to soundscapes.

## 2. Introduction

Certain professions require a high degree of skill and precision but must be performed in environments where distractions are inevitable and mistakes can have fatal consequences. Surgery, for example, is inherently demanding, requiring precise handling of instruments, advanced technological skills, and high levels of concentration over a long duration. In such professions, cognitive resources must be allocated to incoming information across multiple sensory modalities (Wickens, 2008). The environment is characterized by a complex soundscape, comprising a multitude of concurrent sounds, including communication, alarms, monitoring sounds, and instrument usage. The individual team member must distinguish between relevant and irrelevant, potentially distracting, information within this complex soundscape. Thus, distraction represents a cognitive challenge in the operating room (OR), with a notable impact on personnel well-being (Kern et al., 2019) and has been linked to elevated stress levels and an increased likelihood of errors (Mentis, Chellali, Manser, Cao, & Schwaitzberg, 2016).

Although a variety of sounds can be perceived as distracting (Gülşen, Aydıngülü, & Arslan, 2021; Tsiou, Efthymiatos, & Katostaras, 2008; Weigl, Antoniadis, Chiapponi, Bruns, & Sevdalis, 2015), irrelevant speech, that is, speech unrelated to the procedure, is particularly perceived as distracting (Healey, Primus, & Koutantji, 2007; Tsiou et al., 2008; van Harten, Gooszen, Koksma, Niessen, & Abma, 2021) and increases perceived workload (Weigl et al., 2015; Wheelock et al., 2015). Although speech is frequently identified as a distractor in the OR, there is mixed evidence regarding the impact of the OR soundscape, including speech, on surgical performance. Some studies showed performance reductions when the soundscape was compared with silence (e.g., Pluyter, Buzink, Rutkowski, & Jakimowicz, 2010; Siu, Suh, Mukherjee, Oleynikov, & Stergiou, 2010) or when irrelevant speech is the sole distractor (Czerwiec, Vannier, & Courage, 2024). Yet, a realistic OR environment includes multiple overlapping sounds, with silence being a rare condition and irrelevant speech only one of several potential auditory distractors (Gülşen et al., 2021). This highlights the need to study how the combination of irrelevant speech and other overlapping sounds in the OR soundscape influence the individual and affect surgical performance under realistic conditions.

Instead of comparing an OR soundscape or speech with silence, auditory distraction should be investigated in the context of varying task demands. Subjective reports from medical personnel indicate that irrelevant speech is perceived as particularly distracting during phases of high task demand compared to phases of low task demand (Persoon, Broos, Witjes, Hendrikx, & Scherpbier, 2011; van Harten et al., 2021). This suggests that surgical task demand modulates the impact of irrelevant speech distractions. Understanding the contribution of distractors like irrelevant speech and their interaction with task demands is therefore crucial for optimizing performance and improving the work environment.

A comprehensive assessment of the interaction between task demand and distraction in complex environments like the OR requires a combination of objective and subjective measures. Using only subjective measures (e.g., self-reports), it can be challenging to separate the specific contribution of speech from the overall impact of the task demand, and other sounds in the OR environment (Dias, Ngo-Howard, Boskovski, Zenati, & Yule, 2018). Similarly, measures of performance may not capture situations where individuals find the soundscape distracting, even if their performance is unaffected (Rosenkranz, Haupt, Jaeger, Uslar, & Bleichner, 2024). For example, surgeons may experience increased workload in order to maintain a high level of performance within a distracting environment. To complement self-reports and performance measures, electroencephalography (EEG) provides another measurement. Mobile EEG is increasingly employed in the study of work environments (Wascher et al., 2021) and provides reliable responses to complex soundscapes and speech while a task is being performed (Herrmann, 2024; Rosenkranz, Cetin, Uslar, & Bleichner, 2023; Rosenkranz et al., 2024; Xie, Brodbeck, & Chandrasekaran, 2023). Thus, EEG can be related to the different potential auditory distractors like irrelevant speech and the OR soundscape and assess how the processing of each distractor varies with task demand.

To investigate the neural processing of distinct auditory stimuli within the OR sound-scape, we employed two complementary EEG analysis approaches. First, we computed event-related potentials (ERPs) which are well-established for studying the processing of transient sounds in relation to task demands (e.g., Wascher et al., 2021). This makes ERPs particularly well-suited to examine how surgical task demand influences the processing of task-irrelevant auditory stimuli. To account for the continuous aspects of the OR sound-scape, we also computed temporal response functions (TRFs), a robust tool for capturing neural responses to continuous auditory stimuli, including speech and concurrent sounds (Crosse, Di Liberto, Bednar, & Lalor, 2016; Rosenkranz et al., 2023, 2024) By using both ERPs and TRFs, we aim to gain a more comprehensive understanding of how discrete and continuous auditory stimuli are processed and how task demand shapes the neural responses in a complex work-like environment.

The ability to filter out irrelevant auditory information is essential for maintaining focus on a task. Sensory gating, a neural mechanism thought to help suppress responses to repetitive, irrelevant stimuli, may play a key role in this process (Lijffijt et al., 2009). To investigate sensory gating without interfering with the surgical task, we employed the paired-click paradigm. This approach contrasts the neural response to an initial click to a repeated second click. The reduction in response from the first to the second click reflects the strength of sensory gating. Studies suggest that greater cognitive engagement in a task increases the response reduction in the paired-click paradigm, indicating more effective inhibition of irrelevant stimuli (Lijffijt et al., 2009). Similarly, an increase in task demand can suppress the processing of irrelevant auditory stimuli when the relevant stimuli are of a different sensory modality (Molloy, Lavie, & Chait, 2019; Sörqvist, Dahlström, Karlsson, & Rönnberg, 2016; Sörqvist, Stenfelt, & Rönnberg, 2012). Sensory gating effects have been shown to remain robust even when individuals are engaged in cognitive tasks or exposed to background noise (Hölle & Bleichner, 2023; Major et al., 2020), making this method interesting for studying auditory filtering under real-world conditions. Based on these findings, we hypothesized that high task demand would enhance sensory gating, resulting in a larger difference between ERPs in responses to the first and second clicks, compared to low task demand.

Besides examining ERPs in response to the paired-click paradigm, we investigated whether the reduction in ERP amplitudes under high compared to low task demand (Molloy et al., 2019; SanMiguel, Corral, & Escera, 2008; Sörqvist et al., 2016) extends to neural responses to the entire soundscape. Unlike the repetitive stimuli typically used in ERP studies, natural soundscapes consist of distinct and concurrent, as well as, overlapping auditory events. To better capture how such soundscapes are processed, we computed a general neural response using TRFs. This approach allowed us to assess whether the reduction in ERP responses under high task demand could also be observed when analyzing the continuous OR soundscape.

For speech processing, we expected it to be modulated by task demand as well, though the direction of this effect is uncertain. While prior studies indicate that processing of irrelevant non-speech sounds decreases with increasing task demand (SanMiguel et al., 2008; Sörqvist et al., 2016), speech has a higher potential to distract than non-speech sounds (Szalma & Hancock, 2011). This makes speech particularly disruptive for OR personnel, especially during phases of high task demand (van Harten et al., 2021; Weigl et al., 2015; Widmer et al., 2018). Given that distracting non-speech sounds enhance the neural response (Huang & Elhilali, 2020), we considered the possibility that the neural response to speech could be enhanced during high-demand phases, even as non-speech sounds are filtered out.

In summary, this study examined how an OR soundscape including irrelevant speech and varying task demands interact to influence self-reports, surgical task performance, and auditory processing, as measured by EEG. We hypothesized that higher task demand will lead to an increase in perceived workload, while the presence of irrelevant speech will increase distraction ratings. Additionally, we explored the interaction between task demand and speech presence to assess their combined influence on self-reported workload and distraction. As workload is a rather general measure of surgical task demand, we also explored how the tasks and speech affect specific aspects of self-reported demand (Wilson et al., 2011). We further explored the effect of irrelevant speech on surgical task performance. Regarding neural responses, we expected that higher task demand will result in an increased sensory gating, as measured by ERPs. For TRF responses, we hypothesized that increased task demand will lead to a lower neural response to an OR playback. Furthermore, we explored the relationship between task demand and speech processing, as well as the association between neural responses to the OR playback and speech with self-reported workload.

## 3. Method

This study involving human participants was reviewed and approved by Medizinische Ethikkommission, Carl von Ossietzky Universität Oldenburg, Oldenburg (2021-031). The participants provided their written informed consent prior to participating in this study. All participants received monetary reimbursement.

### 3.1. Participants

25 participants were recruited through an online announcement on the University board (age range: 19-34; mean age = 24.84; 14 female; 10 medical students). Eligibility criteria included: self-reported normal hearing, normal or corrected vision, no psychological or neurological condition, right-handedness, and no experience with surgery or surgical simulations. In total five participant were excluded from all analyses involving EEG, due to the following reasons: no data were present due to a recording error (N=1); package loss during recording resulted in timing problems (N=2); connector problems resulted in artifactual data (N=2). Those participants were still included in the self-report and performance analyses.

### 3.2. Paradigm

We employed a within-subject design, featuring a 2 (task: easy vs difficult) x 2 (sound: speech present vs speech absent) factorial structure, where each condition was repeated over three blocks, resulting in a total of 12 blocks (Figure 1). The participants were required to complete the surgical tasks peg transfer and suturing. These tasks were selected as they have been demonstrated to elicit either low or high workload, respectively (Lim et al., 2023; Scerbo, Britt, & Stefanidis, 2017). During all blocks, participants were presented with a task-irrelevant soundscape. In all blocks the soundscape consisted of sounds from an actual OR (i.e., OR playback), and click sounds. Additionally, in half of all blocks, speech was presented. Each block lasted up to six minutes, with the end of the block signaled by the soundscape fading out, indicating that participants should stop the surgical task. Participants were instructed that all auditory stimuli were irrelevant and could be ignored.

**Figure 1.**
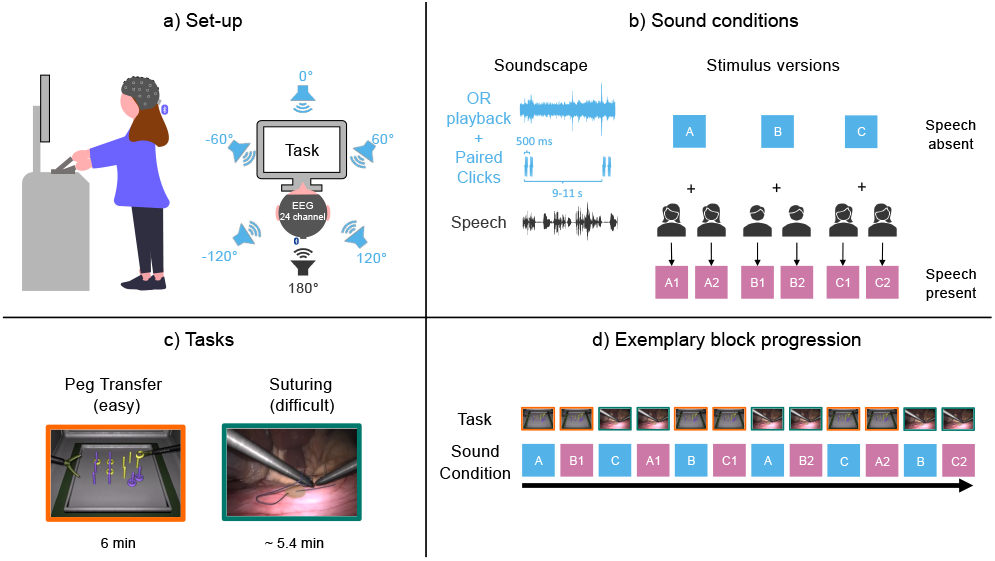
a) Participants performed surgical training tasks while standing in front of a surgical simulator, equipped with a 24-channel mobile EEG cap. A soundscape was presented through a loudspeaker array positioned around them. From five loudspeakers, marked in blue, an OR playback and paired clicks were presented. From one loudspeaker, marked in black, speech was presented. b) All sound conditions (i.e., speech-absent and speech-present) included the OR playback and paired clicks. Additionally, stories from three speaker (two stories per speaker) were presented, with each speaker paired with one of the three OR playbacks. This resulted in six stimuli for the speech-present condition. c) Participants completed two tasks of varying difficulty: peg transfer (easy) and suturing (difficult). The peg transfer task was performed until the end of a block (after six minutes). The suturing task was performed until it was finished (which was on average after 5.4 min), but no longer than six minutes. d) Example block progression for one participant. Each task was presented in two consecutive blocks, alternating between speech-present and speech-absent conditions. The starting task and sound condition was counterbalanced across participants. Additionally, while the overall order of sound conditions was fixed, the starting sound condition was rotated between participants.

#### 3.2.1. Auditory stimuli

We generated nine auditory stimulus versions, each lasting six minutes, which were consistent across all participants (Figure 1b). Of these, three versions featured each a different OR playback with click pairs distributed throughout the playback duration and were used for the speech-absent conditions. The remaining six versions were used for the speech-present conditions, created by pairing each speech-absent version with one of three speakers, with each speaker narrating two different stories. The OR playback and paired clicks were presented through five loudspeakers positioned around the participant (0°, 60°, 120°, -120°, -60°), while speech was presented through a single loudspeaker positioned behind the participant (180°, Figure 1a). Auditory stimuli were presented using Psychtoolbox 3 (v3.0.17, Kleiner et al., 2007). For each stimulus type, a sound marker was generated using the lab streaming layer library (v1.14, Kothe et al., 2024).

##### OR playback

The three OR playbacks were extracted from a recording during a visceral surgery using a field recorder which was positioned close to the surgery table at the University Hospital Oldenburg Rennies et al. (2023). The recording contains a variety of sounds, such as ventilation noise, beeps from monitoring devices, instrument clatter, and instrument sounds. Intelligible speech was removed after the recording for privacy reasons, however, unintelligible muttering and non-vocal sounds such as coughing were preserved.

##### Paired clicks

We presented pairs of clicks, with an interval of 500 ms between clicks and an interval of 9-11 sec between click pairs. Each click pair consisted of two identical clicks (1000 Hz, 4 ms duration, 1 ms onset and offset ramps). In total, 35 pairs were presented per block. To ensure that energetic masking influences of the clicks was similar across participants, all pairs were presented at fixed moments in the OR playback.

##### Speech stimuli

The speech stimuli were chosen from a database containing German speakers, who were telling stories about self-selected content (Daeglau et al., 2023), and have been shown to provide measurable EEG responses (Daeglau et al., 2025; Wiedenmann et al., 2023). The natural speech included speech pauses and filler words which increased the ecological validity of our approach. Three speakers were chosen, each telling two stories. To control for differences in loudness, the speech stimuli were matched to have the same root-mean-square (RMS) value.

#### 3.2.2. Surgical tasks

For the surgical task, the LabSim^®^ (Surgical Science, Sweden) simulator was used. The simulator includes the surgical training tasks peg transfer and suturing which were chosen as they differ in difficulty (Figure 1c; Lim et al., 2023; Scerbo et al., 2017), required bimanual control, and lasted at least three minutes for inexperienced individuals, ensuring that sufficient data could be collected. This minimum duration was an approximation based on observations from pilot data. As the tasks varied in their goal and procedural steps, the performance measures were different between tasks. Peg transfer included the performance measures ‘number of transfers’ and ‘number of drops’, while suturing included the performance measures ‘duration’ and ‘damage’. Although participants were not provided with any feedback regarding their performance during the task, they were instructed as to the performance measures that we investigated. Participants always completed two consecutive blocks of one task (e.g., peg transfer) and then switched to two consecutive blocks of the other task (e.g., suturing).

##### Peg Transfer

In the peg transfer task, participants were required to transfer rings between two pairs of pegs, including a switch between grasping instruments for each transfer. This task was defined as easy, as it involved only few and repetitive procedural steps. Participants were instructed to complete as many transfers as possible within a block with minimal ring drops. The task automatically ended after six minutes. The number of transfers and drops were used as the performance measures; these values were not directly provided by the LabSim software but could be compute based on the number of grasps, average drops, and average transfers.

##### Suturing

In the suturing task, participants were required to drive a needle through tissue and tie two knots in the suture thread using the provided instruments. This task was more difficult than the peg transfer, as it included several procedural steps, and required a higher degree of dexterity. Once the second knot was tied, the task ended and the soundscape stopped. The performance measure were task duration and damage. Damage was defined as the number of times the tissue was touched and the amount of pressure applied to the needle after it was driven through the tissue. This resulted in a damage score from 0 to 100, with a higher score indicating less damage.

#### 3.2.3. Counterbalancing of blocks

To counterbalance the order of tasks, half of the participants started with peg transfer, while the other half started with suturing. The experiment followed a fixed order of stimulus conditions where speech absence and presence alternated with each block (Figure 1d). To counterbalance the sequence of stimulus conditions across participants, participants began at a different starting point within this fixed order and then continued sequentially. Consequently, the order was repeated after the 12th and 24th participant. We further ensured that one story of each speaker was presented during peg transfer and the other story of each speaker during suturing.

#### 3.2.4. Procedure

After arrival, participants practiced the use of the lab simulator for 45 min with the following procedure. To get acquainted to the simulator and the instruments, two simple training tasks (i.e., instrument navigation and grasping) were repeated twice each. This was followed by two blocks of peg transfer and two blocks of suturing. During the first block of each task, the experimenter provided instructions and guidance. During the second block of each task, the experimenter left the room and an OR playback was presented. The OR playback was not used during the experimental blocks. Participants were always allowed to ask questions and watch short instruction videos, provided by the manufacturer of the simulator. After the training, the EEG cap was fitted and participants performed resting measurements (i.e., two minutes of eyes open, two minutes of eyes closed, and listening to a sequence of 20 beeps). After these, participants performed the 12 experimental blocks. After each block participants completed the SURG-TLX (Wilson et al., 2011) thereby providing a self-reported workload measurement. The SURG-TLX contains six items related to the different aspects of surgical demand (mental demand, physical demand, temporal demand, complexity of procedure, stress, and distraction). The items were rated on a visual analogue scale with scores ranging from 0 (low) to 20 (high). At the end of the experiments, participants answered 12 questions regarding the content of the speech stimuli (e.g., Was the topic of one of the stories a bike?). Six questions were related to the content of the stimuli - one question per stimulus - while six questions were unrelated. The participants had to indicate whether they perceived the content or not. An inspection of the speech content questions showed that 86,67% of the questions were answered correctly. Thus, participants could discriminate between speech-related and speech-unrelated questions.

### 3.3. EEG data acquisition

Participants were asked to wash their hair at the day of recording prior to the experiment. EEG data were recorded using a wireless amplifier (SMARTING, mBrainTrain, Belgrade, Serbia) attached to the back of a 24-channel EEG cap (EasyCap GmbH, Hersching, Germany) with Ag/AgCl passive electrodes and the reference and ground electrode at position FCz and AFz, respectively. The data were recorded using a sampling rate of 500 Hz, and transmitted via Bluetooth from the amplifier to a Bluetooth dongle (BlueSoleil) that was plugged into a computer (DELL Precision 3630).

After fitting the cap, the skin at each electrode site was cleaned using 70% alcohol. Skin conductance between the scalp and electrodes was increased using abrasive gel (Abralyt HiCl, Easycap GmbH, Germany). Impedance were kept below 10 kΩ at the beginning of the recording.

The transmitted EEG data and sound marker were recorded in the Lab Recorder software ^1^ and saved as .xdf files. The same computer was used for data recording and sound presentation. Due to technical reasons, a constant delay between the marker indicating sound onset and the actual sound presentation was measured in advance. To account for this delay, the marker was adjusted offline by shifting it 40 ms.

### 3.4. EEG preprocessing

The EEG data were analyzed using EEGLAB (v2022.1, Delorme & Makeig, 2004) in MATLAB R2020b (The MathWorks, Natick, MA, United States). As a first step, bad channels that were recognized during recording were removed, resulting in the removal of one channel for two participants. After bad channel removal, the data were cleaned from artifacts using infomax independent component analysis (ICA). To improve ICA, the data were first high-pass filtered (passband edge = 1 Hz^1^), low-pass filtered (passband edge = 40 Hz^2^), and resampled to 250 Hz. Then, the resting data and data of each block during which audio was presented, were cut into consecutive epochs of one second. To minimize artifacts from the start and end of the task the first and last five seconds of each block were excluded. Improbable epochs with a global (all channels) or local (single channel) threshold exceeding 5 standard deviations were automatically rejected using the *jointprob* function. ICA decomposition was applied to the remaining epochs. The resulting components were back-projected on the raw data. The raw data were then high-pass filtered (passband edge = 1 Hz^1^) and low-pass filtered (passband edge = 25 Hz^3^). The back-projected components were then classified using the EEGLAB toolbox *ICLabel* (Pion-Tonachini, Kreutz-Delgado, & Makeig, 2019) with the ‘lite’ classifier which is better at detecting muscle artifacts than the default classifier (Klug & Gramann, 2021). Components belonging to the categories eye blink and movement or muscle movement with 70% confidence or heart with 80% confidence were removed. Note, that the ICLabel classifier did not classify all components correctly because it was trained on stationary data with a larger electrode setup than ours. Therefore, we manually checked the components and made the following adjustments: Components indicating lateral eye-movement were not always correctly classified and individually removed. Furthermore, channel Tp9 and Tp10 contained noise from muscle movement. We observed this already in a previous experiment, where a surgical simulator was used (Rosenkranz et al., 2024). We assume that the bi-manual control of the simulator activates neck muscles, resulting in artifacts in electrodes that are close to the neck. As channel Tp9 and Tp10 are used for re-referencing, we removed components showing muscle activity in Tp9 and Tp10. Afterwards, previously rejected channels were interpolated using spherical interpolation. Lastly, channels were re-referenced to the linked mastoids (Tp9 and Tp10).

### 3.5. ERP analysis

We analyzed the neural response to the irrelevant soundscape using two different approaches. Event-related potentials (ERPs) were computed in response to the paired clicks, while a temporal response function (TRF) was used to analyze the neural response to the OR playback and speech. Since the irrelevant speech was presented separately to the OR playback, we computed separate TRFs for these.

We quantified ERP amplitudes using the following procedure: For each block we extracted epochs from -200 to 1000 ms relative to the onset of the first click and baseline-corrected the epochs from -200 to 0 ms. Epochs exceeding a threshold of 3 standard deviations globally (across all channels) or locally (within a single channel) were automatically rejected using the *jointprob* function. Data from channel FC1, FC2, Fz, and Cz were then averaged, as the auditory N1 and P2 ERP components are prominent at these channels (Crowley & Colrain, 2004; Näätänen & Picton, 1987). To extract the N1 and P2 components for both the first and second clicks, we first computed an average ERP across participants for each sound condition. It should be noted that the extraction of time windows was conducted for the two sound conditions separately to account for the acoustic differences between the two conditions. For each sound condition, we identified the N1 peak within the range of 80 to 140 ms. Amplitudes were then averaged within a *±* 25 ms range around the peak, resulting in one N1 amplitude value per participant and task for the first click. The P2 component was identified similarly, with a peak search window of 150 to 250 ms and a *±* 25 ms range around the peak. For the second click, we added 500 ms to the N1 and P2 time-window of the first click, and averaged across this time-window. We performed a peak-to-peak analysis by subtracting the N1 amplitude from the P2 amplitude for each click. The resulting difference score defined the response amplitude for each click, sound condition, task, and participant. The difference between the first and second click defined the gating value.

### 3.6. TRF analysis

#### 3.6.1. Audio preprocessing

To relate the ongoing soundscape to the ongoing neural response, we extracted the envelope of the OR playback and speech. The OR playback included noise from running machines and ventilation, which produced an envelope with little variation. Such low variability in the envelope can lead to poor TRF estimation (Rosenkranz et al., 2023). To address this, we applied a Wiener filter implemented in MATLAB (Plapous, Marro, & Scalart, 2006; Scalart, 2023). For this, we first high-pass filtered each OR playback at 1 Hz^4^. We then estimated the power spectral density of the noise using the first second of each OR playback, as it was representative of the static noise in the OR playback. The noise estimate was then subtracted from the remaining signal. Afterwards, we extracted the envelope from the noise-reduced OR playbacks and raw speech using the *mTRFenvelope* function Crosse et al. (2016, 2021). We resample all envelopes to 125 Hz.

#### 3.6.2. Backward modeling

A backward modeling approach was utilized to analyze the response to each stimulus envelope separately using the mTRF toolbox (Crosse et al., 2016, 2021). In backward modeling, the neural response is used to reconstruct features of the stimulus. The procedure involved a nested cross-validation for each task and stimulus. As a first step, the EEG data were resampled to 125 Hz to match the sampling rate of the envelopes. The first and last five seconds of the EEG data and envelope of each block was removed. Each block was split in half, resulting in six segments per task. The mean duration of each segment was 175 and 162 seconds for the peg transfer and suturing task, respectively. Each segment served once as a test set while the remaining segments served as training sets, resulting in six folds. For each fold, the following procedure was employed: The training segments were cross-validated to determine the optimal regularization parameter (i.e., lambda) using the function *mTRFcrossval*. The optimal lambda was searched for in the range of 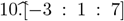. A time-window from 0-300 ms was chosen, as irrelevant stimulus processing typically takes place in earlier time-windows (Hausfeld, Riecke, Valente, & Formisano, 2018; Hausfeld, Shiell, Formisano, & Riecke, 2021). The lambda corresponding to the maximum correlation coefficient was selected for model training. The model was then trained using the *mTRFtrain* function and the optimal lambda. The trained model was then applied to the test set using the *mTRFpredict* function, which correlated the actual and predicted stimulus envelope. This resulted in one Pearson correlation value (i.e., prediction accuracy) per fold which were averaged across folds, resulting in one correlation value per participant, task, and stimulus. In all conditions, we analyzed the neural response to the OR playback. Additionally, in the speech-present condition, we applied the same analysis procedure to the speech stimuli. For this, we computed TRFs using the combined speech material, disregarding differences in content, speaker sex, and speech characteristics, such as word frequency.

We also investigated whether the correlation between the actual and reconstructed stimulus envelope was above chance. For this, a permutation test was conducted using the *mTRFpermute* function. This involved shuffling the response data and recalculating the correlation to generate a distribution of correlation coefficients under the null hypothesis, representing chance-level performance. In total 17 permutations were performed per fold, resulting in 102 permutations per participant, task, and stimulus. The permutation test of each fold used the optimal lambda of the respective fold. The chance-level was defined as the 95th percentile across all permutations of a stimulus.

### 3.7. Statistical analyses

All statistical analyses were performed in R Studio (v4.2.1). For most outcome measures, we fitted a series of linear mixed models using the R packages lmer4 (v1.1-30) (Bates, Mächler, Bolker, & Walker, 2015). We subsequently added fixed then random effects and evaluated the improvement in models fit. The baseline model always included the random intercept of the participant. For most models, the predictors of *task* and/or *sound* were added. *Task* contained two categories, peg transfer and suturing, which were coded 0 and 1, respectively. *Sound* contained two categories, speech absent and speech present, which were coded 0 and 1, respectively. The best fitting model was determined using likelihood-ratio testing. We report results from the likelihood-ratio test comparing a model with a fixed or random effect to a model without the effect. We further report for the fixed effects the *b* value and standard error (SE) of the best fitting model. The model comparisons for all computed models can be found in the supplementary materials (section ‘Model comparisons’).

#### 3.7.1. Self-reported workload

For self-reported workload and distraction we used two outcome measures from the SURG-TLX. First, the total SURG-TLX score defined the workload and was calculated by averaging the scores for all items. For each task and sound condition, the mean score was calculated across blocks. Secondly, the score for the item distraction was calculated by averaging across blocks. This resulted in 2 (task: peg transfer vs suturing) x 2 (sound: speech present vs. speech absent) workload and distraction scores for each participant. We expected the total score to change between the tasks, and explored the effect of the sound condition. Therefore, we iteratively added first *task*, second *sound*, and then their interaction as fixed effects. After fitting the fixed effects, we fitted the random effects by adding *task* and *sound* as random slopes.

For the distraction item we expected it to differ mainly between sound conditions, thus we first added *sound*, second *task*, and then their interaction. After fitting the fixed effects, we fitted the random effects by adding *sound* and *task* as random slopes.

While the total score and distraction item were our main outcome measure, we also explored how *task* and *sound* affect the different aspects of surgical demand. Therefore, we followed the same procedure as for the total score also for the other items of the SURG-TLX (i.e., mental demand, physical demand, temporal demand, complexity, and stress).

#### 3.7.2. Surgical task Performance

We computed separate linear mixed models for the outcome measures of the peg transfer and suturing tasks. As the outcome parameters were different for each task, we did not investigate whether performance differed across tasks. Instead, we focused on whether the presence of speech influenced task performance. For each outcome parameter, we investigated the effect of adding the fixed and random slope of *sound*. For the outcome measure ‘duration’ the data were skewed, as most participants did not complete the task within the 6 minutes. Therefore, we computed a beta mixed model (Verkuilen & Smithson, 2012) using the R package *glmmTMB* (v.1.1.7). For this, we normalized all values between 0 and 1 with 1 indicating that the entire duration was used. As beta models do not allow values to be exactly 1 we transformed boundary values by subtracting 0.002. Otherwise, the same method as with the other outcome parameters was applied.

#### 3.7.3. Sensory Gating

We use the paired-click paradigm to assess the amount of sensory gating for each task. We first evaluated whether a gating effect was present. We did this separately for the sound conditions, as they were acoustically different, thus different responses could be expected. For each sound condition we evaluated whether the response to the first and second click were different, in other words, whether gating was present. We computed a linear mixed model for each sound condition using the response amplitudes to the clicks as an outcome measure. The baseline model included *participant* as a random intercept and *position* (coded 0 and 1 for the response to the first and second click, respectively) was subsequently added as a fixed effect and random slope.

We then checked, whether gating differed between tasks. The gating value was obtained by subtracting the amplitude of the first click from that of the second click. This gating value, calculated for each participant and task, reflects the strength of the sensory gating, with a larger value indicating a larger gating, in other words, better suppression of the second click after hearing the first one. For each sound condition, we computed the baseline model including *participant* as a random intercept and subsequently added *task* as a fixed effect and random slope.

#### 3.7.4. Continuous stimuli

For the response to the continuous stimuli, three prediction accuracies were computed. For the speech-absent condition, only the playback stimulus was modeled, resulting in one prediction accuracy. For the speech-present condition, the playback and speech were modeled individually, resulting in two prediction accuracies. Each prediction accuracy was modeled using the baseline model *participant* as a random intercept and subsequently adding *task* as a fixed effect and random slope.

#### 3.7.5. Exploratory analyses

We investigated the relationship between self-report and neural measurements. To do this, we used the prediction accuracies (i.e., *r*) as continuous predictors for the total workload score. We calculated the average workload score for both the blocks where speech was absent or present. In both the speech-absent and speech-present conditions, the prediction accuracy of the OR playback was used to predict the total workload for each respective condition. In addition, in the speech-present condition, the prediction accuracy of the speech stimulus was also used to predict the total workload for that condition. To receive meaningful *b* estimates we centered the prediction accuracies for each stimulus. To account for the effect of task, *task* was also included as a predictor. Thus, the baseline model included the random effect *participant* and fixed effect *task*. We subsequently added the fixed effect *r* and the interaction between *r* and *task*. This was done for each stimulus separately.

## 4. Results

### 4.1. Self-reported workload

**Total score** Most participants reported a higher overall workload during the suturing compared to peg transfer task (Figure 2a). The best fitting model included the main effects *task* (*χ*^2^(1) = 85.86, *p <* .001, *b* = 4.58, *SE* = 0.5) and *sound* (*χ*^2^(1) = 12.01, *p <* .001, *b* = 1.2, *SE* = 0.27), but no interaction (*χ*^2^(1) = 0.0001, *p* = .993). When allowing the effect of *task* and *sound* to vary across participants, the model fit further improved (*task* : *χ*^2^(1) = 24.68, *p <* .001; *sound* : *χ*^2^(1) = 10.26, *p* = .016). This indicates that participants experienced higher self-reported workload during the suturing task compared to the peg transfer task, a higher workload when speech was present compared to when speech was absent, and that the strength of both effects varied between participants.

**Figure 2.**
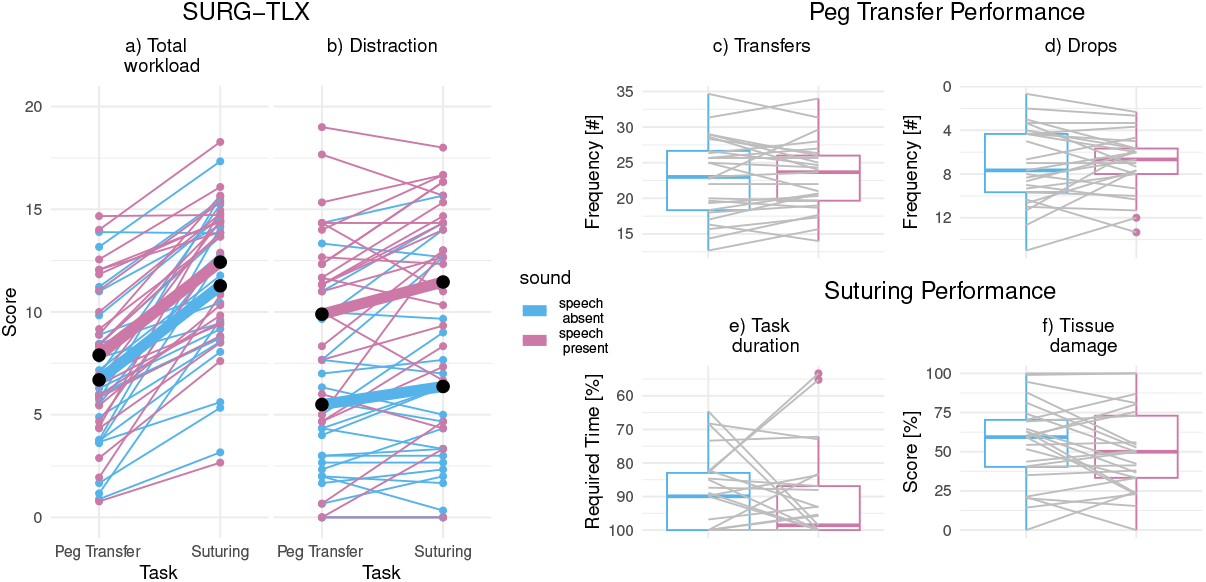
Self-report and performance measures for each task and sound condition. The tasks had two difficulty levels, the easy peg transfer task and the difficult suturing task. Self-reports were derived from the SURG-TLX. a) The total workload score (task & sound: *p <* .001). b) The score on the distraction item (task & sound: *p <* .01). Thin lines represent individual data for each sound condition, thick lines the average across participants. To evaluate the effect of speech on surgical task performance we used for the peg transfer task c) the number of transferred ring, and d) the number of dropped rings. For the suturing task we used e) the amount of the total time that was required (i.e., 0-6 min, Mean = 5.4 min), and f) the efficacy of handling the tissue. For all performance plots, a high y-axis value indicates good performance, and a low value bad performance. The gray lines show individual participants. None of the performance measures was significantly affected by the sound condition.

#### Distraction score

The best fitting model included the main effects *sound* (*χ*^2^(1) = 60.30, *p <* .001, *b* = 4.79, *SE* = 0.69) and *task* (*χ*^2^(1) = 6.78, *p* = .009, *b* = 1.27, *SE* = 0.33), but no interaction (*χ*^2^(1) = 0.66, *p* = .415). Allowing the effect of *sound*, but not of *task*, to vary across participants further improved the model fit (*sound* : *χ*^2^(2) = 19.37, *p <* .001, *task* : *χ*^2^(3) = 2.30, *p* = .513). This indicates that participants experienced higher levels of distraction during the suturing task compared to the peg transfer task, more distraction in the presence of speech compared to its absence, and that the effect of *sound* varied between participants.

#### Exploratory analyses of individual SURG-TLX items

We investigated the remaining SURG-TLX items individually (see supplementary figure S1) and listed the results in the supplementary table S1. To summarize, we subsequently added the fixed effects, *task, sound*, and their interaction, and the random slopes. Adding *task* as fixed effect as well as random slope significantly improved model fit for the items mental demands, physical demands, temporal demands, and complexity of procedure. For the same items, adding *sound* or the interaction between *task* and *sound* did not improve the model fit significantly. For the item situational stress, the best fitting model included the main effects for *task* and *sound* and random slopes for both effects. To summarize, while all items showed higher scores for the suturing compared to peg transfer task, only the distraction and stress item showed also higher scores for the speech-present compared to speech-absent condition.

### 4.2. Surgical task performance

For the peg transfer task, the outcome parameters were the number of transfers and the number of drops. For the suturing task, the outcome parameters were the time required to complete the task and the extent of damage. As shown in Figure 2 c)-f), adding *sound* as predictor did not change model performance for any outcome measure (Transfers: *χ*^2^(1) = 0.021, *p* = 0.886; Drops: *χ*^2^(1) = 0.127, *p* = 0.7215; Time: *χ*^2^(1) = 0.22, *p* = 0.64; Damage: *χ*^2^(1) = 2.267, *p* = 0.132). This indicates that the presence of speech did not change the measured task performance during either surgical task. Furthermore, there was a high variability in performance between participant. This was probably the result of our participant pool, that had little laparoscopic experience.

### 4.3. Sensory gating

To investigate the presence of sensory gating, we computed whether the ERP amplitude changed from the first to the second click. We observed a gating effect in the speech-present and speech-absent condition. For both sound conditions, the model fit improved when adding *position* as a predictor (Figure 3b & e; speech-absent condition: *χ*^2^(1) = 29.07, *p <* .001, *b* = *−* 1.1, *SE* = 0.21; speech-present condition: *χ*^2^(1) = 7.81, *p* = .005, *b* = *−* 0.6, *SE* = 0.21). Allowing *position* to vary across participants improved the model fit for the speech-absent condition (*χ*^2^(1) = 11.02, *p* = .004) but not for the speech-present condition (*χ*^2^(1) = 2.67, *p* = .263). Figure 3a) and d) shows that the ERP morphologies between the sound conditions are different, likely due to the different acoustics in each sound condition (i.e. the presence or absence of speech).

**Figure 3.**
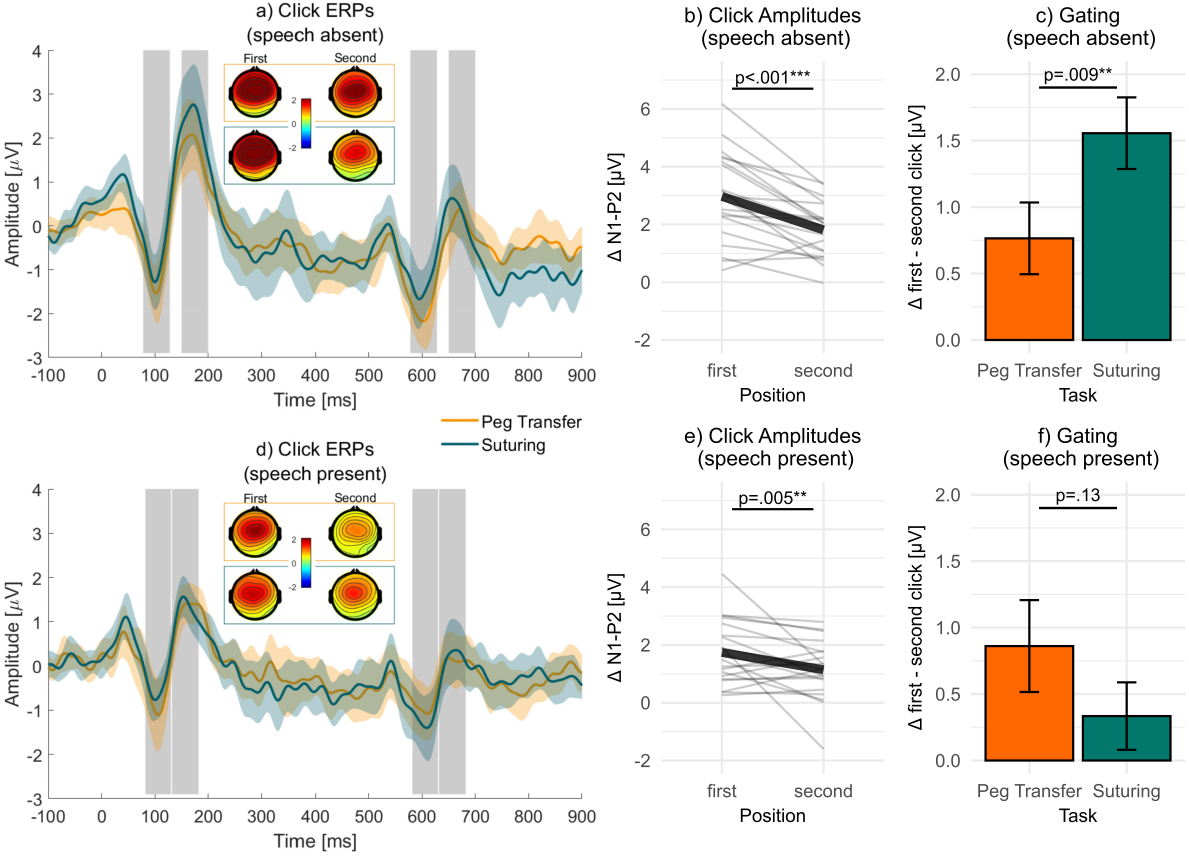
ERPs in response to the paired-click paradigm. The top and bottom row show data for the speech-absent and speech-present condition, respectively. The first column (a&d) shows the ERP time-course and topographies in response to the clicks, which were presented at 0 and 500 ms. The time-course shows the averaged data across participants and channels FC1, FC2, Fz, and Cz (solid line), along with the confidence interval (shaded area) for each task. The gray areas highlight the N1 and P2 time windows. The topographies show the peak-to-peak difference from the averaged amplitudes in the N1 and P2 time-window for the first and second click. The second column (b&e) shows the averaged amplitude for the first and second click. The thick line shows the average across participants and thin lines individual participants. The third column (c&f) shows the strength of gating, that is the difference between the response to the first and second click for each task (*±*1*SE*).

We then investigated whether the sensory gating strength changed between tasks. For the speech-absent condition, adding *task* as a predictor significantly improved model fit (Figure 3c; *χ*^2^(1) = 6.75, *p* = .009, *b* = 0.791, *SE* = 0.287), but adding *task* as random slope led to an unidentifiable model. For the speech-present condition, adding *task* as a predictor did not improve model fit, compared to the baseline model (Figure 3f; *χ*^2^(1) = 2.4, *p* = .13). To summarize, sensory gating was stronger in the suturing compared to peg transfer task in the speech-absent condition.

### 4.4. Responses to continuous stimuli

Figure 4 (top row) shows that all models performed above chance. For the OR playback, adding *task* as a predictor neither increased model performance when speech was absent (*χ*^2^(1) = 1.56, *p* = 0.135) nor when speech was present (*χ*^2^(1) = 0.625, *p* = 0.54). For the speech stimulus, adding *task* as a predictor improved model performance (*χ*^2^(1) = 11.723, *p <* 0.001, *b* = *−* 0.015, *SE* = 0.004), but adding *task* as random slope led to an unidentifiable model. This indicates that the prediction accuracy for speech was higher for the peg transfer than for the suturing task.

**Figure 4.**
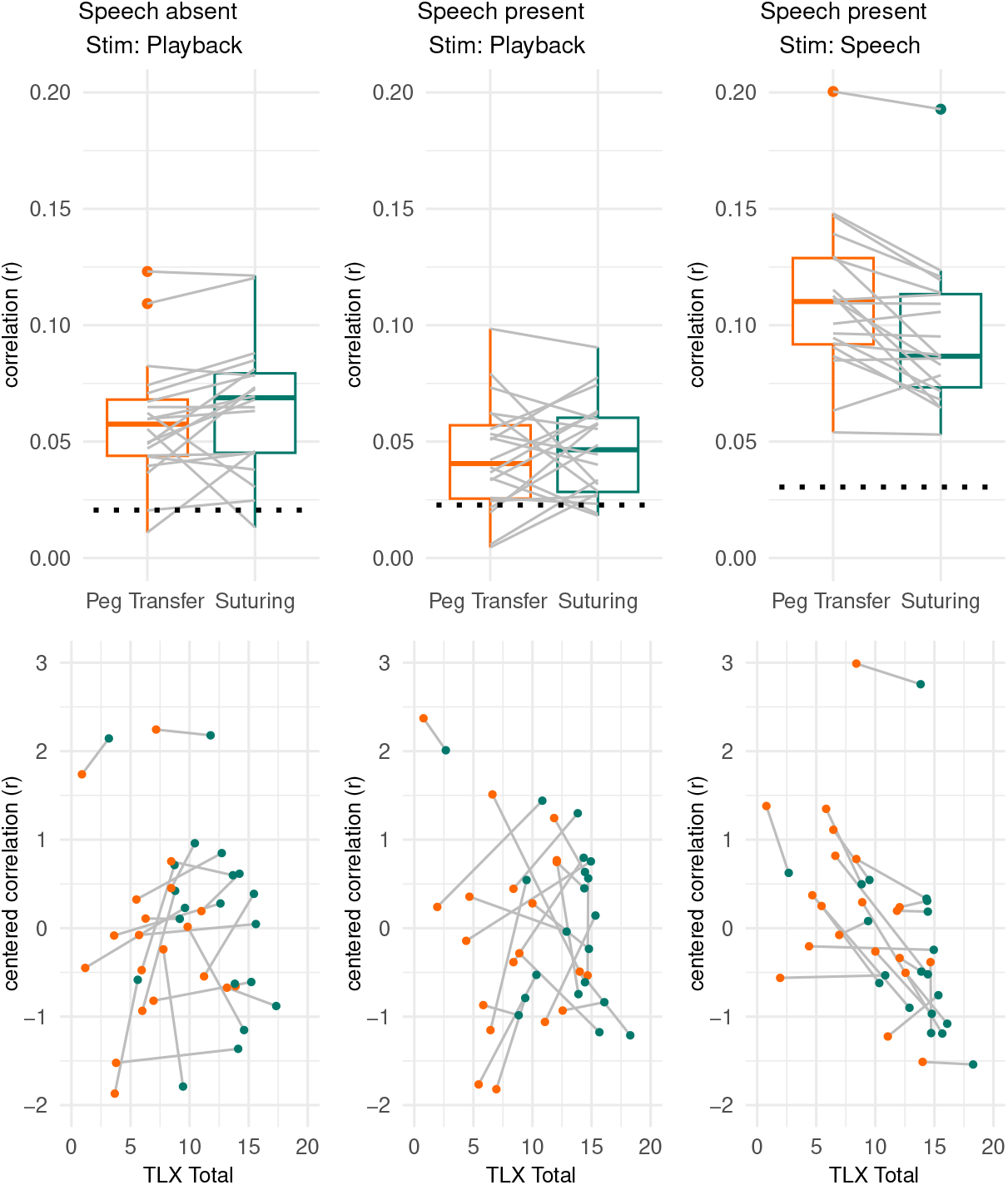
The plots show the effect of task on the correlation between the predicted and reconstructed stimulus (top row) and the effect of task and centered correlation on the total workload score (bottom row). Each column represents reconstruction accuracies for one of the three stimuli (i.e., Stim). Grey lines show data for individual participants. For the top row, the chance level is represented by the black dotted line. For the bottom row, we predicted the workload score using the centered correlation, but plotted the workload score on the x-axis and centered correlation on the y-axis, to visually match the top row.

### 4.5. Relationship between self-report and neural responses

We explored the relationship between the self-report workload measures and neural responses to the continuous stimuli (Figure 4 bottom row). Adding *r* as predictor did not change model performance for the OR playback when speech was absent (*χ*^2^(1) = 0.803, *p* = 0.37) nor when speech was present (*χ*^2^(1) = 0.738, *p* = 0.39). However, adding *r* as predictor significantly improved model performance for speech (*χ*^2^(1) = 4.687, *p* = 0.03, *b* = *−* 1.39, *SE* = 0.64), indicating that a lower neural response to speech was associated with larger perceived workload. Adding the interaction of *task* and *r* did not improve the model further (*χ*^2^(1) = 0.041, *p* = 0.84), and adding *task* or *r* as a random factor lead to unidentifiable models.

## Discussion

While speech distraction in the OR is an often reported problem, only few studies investigated this experimentally. Therefore, we studied how a soundscape, consisting of an OR playback, paired clicks, and irrelevant speech is perceived and processed during the performance of an easy and difficult surgical task, namely peg transfer and suturing, respectively. To understand how the soundscape is processed and influences the individual, we employed self-report, performance, and neurophysiological measurements.

As expected participants reported higher workload for the suturing task compared to the peg transfer task. This aligns with previous research showing that suturing (i.e., the difficult task) to be more demanding than peg transfer (i.e., the easy task), thus resulting in higher workload (Lim et al., 2023; Scerbo et al., 2017). The presence of speech increased the perceived workload as reflected in SURG-TLX items *distraction* and *stress*. Speech likely introduced a salient distraction that was difficult to ignore. This parallels findings from the actual OR where disturbance due to irrelevant speech correlates most strongly with these two items (Weigl et al., 2015). This consistency highlights the realism of our setup in simulating speech distraction within the OR environment, and shows that irrelevant speech increases the overall perceived workload by increasing distraction and stress.

Participants reported feeling more distracted by their environment during the difficult task. This replicates a finding from our previous study where participants perceived the soundscape as distracting primarily when task demand was high (Rosenkranz et al., 2024). This effect is also consistent with reports from OR personnel, who often cite noise as particularly distracting during high-workload phases (van Harten et al., 2021). To address this, noise interventions should prioritize reducing unnecessary noise, especially during periods identified by surgeons as highly demanding. As potential solutions, one could consider either implementing a system to signal that the room should remain quiet, such as a traffic light system (e.g., Engelmann, Neis, Kirschbaum, Grote, & Ure, 2014) or enabling surgeons to reduce potentially unavoidable, task-irrelevant sounds, by using hearing devices (e.g., Leitsmann et al., 2021; Rennies et al., 2023).

While participants’ self-reports indicate more distraction when speech was present, their surgical task performance remained unaffected by speech. Similarly, noise reduction interventions in the OR may increase subjective well-being, without necessarily impacting patient outcomes (Engelmann et al., 2014; Leitsmann et al., 2021). We propose two potential explanations for this discrepancy. First, the maintenance of performance in the presence of task-irrelevant speech comes at the cost of increased workload. This compensatory workload may have long-term implications for surgeons’ well-being, even if patients are not immediately affected (Ayas, Donmez, Kazlovich, Lombardi, & Jain, 2022). Second, performance measurements may not be sensitive enough to detect subtle differences in behavior. For instance, expert and novice surgeons can achieve similar performance on simple surgical tasks, even when distracted (Hsu, Man, Gizicki, Feldman, & Fried, 2008). Furthermore, the evidence regarding the effects of noise on surgical performance are rather mixed (Mentis et al., 2016), which may be caused by the heterogeneity in performance measures across studies. This highlights the limitations of current performance metrics which may only decline under severely distracting conditions, for instance when several distractors are combined (Pluyter et al., 2010; Szafranski, Kahol, Ghaemmaghami, Smith, & Ferrara, 2009). Overall, our results suggest that while the measured performance may not suffer, irrelevant speech adds a mental strain that is reflected in self-reports rather than in surgical performance. This suggests that improving the acoustic environment can benefit surgeons by reducing perceived distraction and workload, contributing to their overall well-being.

As the recording of self-reports is limited to a single point in time, and performance may not accurately reflect changes in the processing of the soundscape, we utilized EEG to objectively examine responses to the soundscape. We employed two approaches, namely ERPs and TRFs to study responses to the transient clicks and continuous stimuli, respectively. With regard to ERPs, a gating effect was observed in both sound conditions, with participants showing reduced neural responses to the second click compared to the first. This replicates previous findings of sensory gating in complex environments (Hölle & Bleichner, 2023; Major et al., 2020). Sensory gating is a neural mechanism that reflects early processing, specifically the filtering of irrelevant information (Lijffijt et al., 2009). Thus, our results show that participants were generally able to effectively filter out repetitive, irrelevant sounds.

While sensory gating was present in both sound conditions, we found that gating strength was affected by task demand when speech was absent, but found no effect when speech was present. In the speech-absent condition, the strength of gating was stronger during the more difficult task, suggesting enhanced suppression of irrelevant auditory stimuli under high task demands. Our findings align with research showing enhanced filtering of irrelevant stimuli at an early stage of processing when task demand was high (Bidet-Caulet, Bottemanne, Fonteneau, Giard, & Bertrand, 2015; Miller, Rietschel, McDonald, & Hatfield, 2011; Sörqvist et al., 2016, 2012). The increased demands of the more difficult task appear to have strengthened sensory filtering, thereby protecting cognitive functioning from the irrelevant stimuli (Lijffijt et al., 2009). Importantly, the paired-click paradigm does not rely on attention markers, such as the P3, which are commonly used in beyond-the-lab studies (Grasso-Cladera, Bremer, Ladouce, & Parada, 2024). Thus, exploring this mechanism across different tasks provides a promising opportunity to investigate how irrelevant sounds are processed without interfering with concurrent tasks (Hölle & Bleichner, 2023).

When speech was present, we found no effect of task demand on gating strength. One possible explanation is that the speech may have masked the click sounds through energetic masking, where the loudness of the speech obscured the clicks (Brungart, Simpson, Ericson, & Scott, 2001; Hölle & Bleichner, 2023; Shinn-Cunningham, 2008). This masking likely made the clicks less perceptible, reducing their processing and leading to a lower signal-to-noise ratio (SNR) for the ERPs. This is evident in the descriptively lower ERP amplitudes for each click in the speech-present condition compared to the speech-absent condition. Consequently, the reduced SNR may have decreased the likelihood of finding an effect, as reflected in the weaker gating effect observed in the speech-present compared to speech-absent condition. Alternatively, the speech may have engaged more perceptual resources than the clicks. Stimuli, that are perceived as separate auditory objects, characterized by variations in spatial location or specto-temporal content (Griffiths & Warren, 2004), are also processes separately (Hausfeld et al., 2018). The separation may lead to a preference to process speech, thereby allocating less resources to the processing of the clicks (Shinn-Cunningham, 2008). This in turn limited the gating effect. Future research could explore these explanations by investigating the influence of irrelevant speech on the processing of other irrelevant sounds.

Utilizing the paired-click paradigm, we investigated how the processing of auditory stimuli was affected by task demand. However, reliance on artificially induced stimuli remains a limitation when transitioning to real-world research (Grasso-Cladera et al., 2024; Matusz, Dikker, Huth, & Perrodin, 2019; Wascher et al., 2021). Therefore, we also investigated the processing of the more realistic parts of the soundscape, incorporating the OR playback and speech. Neural responses to irrelevant speech were reduced during high task demand, suggesting that increased workload leave fewer resources available to process speech. Our findings align with previous studies that demonstrated diminished processing of irrelevant stimuli when task demand increased (SanMiguel et al., 2008; Sörqvist et al., 2016, 2012). These studies were conceptually similar to our, as they investigated irrelevant stimuli with a different modality than the task-stimuli. However, these studies used simple tasks and discrete stimuli, such as single tones. Our study extended this work by demonstrating similar effects with continuous and naturalistic stimuli, namely spoken stories. This provides evidence that such mechanisms persist in realistic scenarios. Furthermore, the prolonged allocation of limited cognitive resources to a demanding task while simultaneously suppressing distracting stimuli can be exhausting (Esterman & Rothlein, 2019). This effort is likely to increase when the target and distractor stimuli share the same sensory modality, as competition for cognitive resources increases (Wickens, 2008). For instance, task-relevant speech becomes more difficult to comprehend when presented alongside an OR playback (Way et al., 2013). Investigating how relevant and irrelevant speech compete for cognitive resources while performing another non-auditory task represents a critical step towards understanding distraction in realistic OR scenarios. Previous research has shown larger neural responses to relevant compared to irrelevant speech during a walking task (Straetmans, Adiloglu, & Debener, 2024; Straetmans, Holtze, Debener, Jaeger, & Mirkovic, 2021). Extending such investigations to surgical tasks of varying difficulty would provide insights into the interplay between dual-tasking (i.e., performing a task while processing speech) and speech distraction in high-demand environments like the OR.

The speech suppression effect emerged at an early stage of processing, as indicated by the time-lag of 0 to 300 ms used in the analysis of continuous stimuli. This is similar to the speech literature that usually employ dual- or multi-talker paradigms, showing that ignored speech processing occurs early (Hausfeld et al., 2018, 2021). This also aligns with the impact of task difficulty on the early ERP components of the paired-click paradigm and suggests that irrelevant speech filtering starts at an early processing stage.

Our exploratory analyses further revealed an inverse relationship between neural responses to speech and self-reported workload, even when accounting for task difficulty. In other words, the strength of the neural response to speech reflected the amount of workload participants experienced. While neurophysiological measures are increasingly used to assess workload, they often rely on ERPs elicited by artificial and repetitive sounds, which are rarely encountered in real-world environments (Wascher et al., 2021). We demonstrated that neural responses to naturalistic stimuli, such as speech, can also serve as markers of perceived workload. This finding could encourage future studies investigating complex work environments to use natural stimuli.

For the OR playback, we found above chance correlations for most participants, showing that stimuli such as an OR playback can be reconstructed from neural data. Contrary to our hypothesis, we found no task modulation for responses to the irrelevant OR playback. While we found a modulated ERP to transient, irrelevant sounds, this did not generalize to all irrelevant sounds. We cannot rule out the possibility that certain sounds within the playback received modulations that were not captured in the computation of a general response to the OR playback. One explanation for this is a discrepancy between our definition of the stimulus categories in our analyses and the actual perception of the OR playback. We defined the clicks, irrelevant speech, and OR playback as separate auditory objects (Griffiths & Warren, 2004). This definition may be applicable to both clicks and speech, as each stimulus exhibited comparable sound characteristics, including amplitude and frequency, over time (Shinn-Cunningham, 2008). However, the OR playback may have been perceived as containing many different auditory objects where each object may have resulted in a different response and task modulation (Huang & Elhilali, 2020). By conceptualizing the entire OR playback as a single auditory object, we may have overlooked sounds within the OR playback that exhibited comparable top-down modulations as observed with the clicks and speech. An alternative explanation is that our analyses may have been biased to enhance speech responses. To have comparable results between the OR playback and speech, we employed the envelope as a feature for both stimuli. This feature is commonly used to maximize speech responses (Crosse et al., 2016). While the envelope is also suitable for describing continuous stimuli beyond speech (e.g., Di Liberto, Pelofi, Shamma, & de Cheveigné, 2020; Hausfeld et al., 2018; Rosenkranz et al., 2023, 2024), alternative representations, such as the mel-spectogram, explain more variance of the neural data (Di Liberto, O’Sullivan, & Lalor, 2015; Haupt, Rosenkranz, & Bleichner, 2025). Such representations may therefore be more sensitive to detect attention modulations. Either way, when dealing with soundscapes that contain multiple overlapping sounds, a more detailed description, for example by classifying sounds according to their content or acoustic features, may further improve the analyses of responses to the soundscape.

Whilst the present study systematically examined the processing of irrelevant speech and its effect on task performance, certain limitations should be noted. The short duration of the tasks, the performance metrics used, and the relatively high variability of performance in the sample of inexperienced participants may have limited the ability to detect subtle performance impacts (Mentis et al., 2016). Furthermore, the ability to react to distractors is likely to change with experience (Hsu et al., 2008). As the present study focused on participants with no prior experience of the OR, the results may reflect more general demand modulations on auditory processing that are susceptible to change with experience. Hence, future studies utilizing longer tasks and experienced personnel could offer further insights into the impact of speech distraction on surgical personnel.

In the strive to understand auditory distraction in a real-world setting, we combined self-reports, performance, and neurophysiological measures. We investigated how a complex soundscape, consisting of an OR playback, paired clicks, and irrelevant speech is perceived and processed during the performance of surgical tasks that varied in difficulty. Our study demonstrates that while irrelevant speech may not immediately impact performance, it increases perceived workload and distraction during the performance of difficult tasks. This finding could be generalized to other high-stakes settings where a control of the auditory environment is necessary to support the well-being of personnel. Furthermore, our study adds to the growing body of literature that utilizes EEG beyond the lab and at workplaces (e.g., Dehais, Karwowski, & Ayaz, 2020; Grasso-Cladera et al., 2024; Wascher et al., 2021). Using mobile EEG we could investigate the underlying neurophysiological mechanisms of processing the irrelevant aspects of the soundscape, thereby avoiding a direct interference with the surgical task. This showed that irrelevant speech responses were reduced at an early stage of stimulus processing when performing a difficult task. Furthermore, the response was inversely related to workload. This highlights the potential of using mobile EEG to investigate workload in complex, real-world settings with naturalistic stimuli, paving the way for more effective strategies to monitor and mitigate auditory distraction in high-stakes environments.

## Footnotes

1 https://github.com/labstreaminglayer/App-LabRecorder, v1.14

1a filter order = 825, transition bandwidth = 1 Hz, cutoff frequency (−6dB) = 0.5 Hz

2 filter order = 83.5, transition bandwidth = 10 Hz, cutoff frequency (−6dB) = 45 Hz

3 filter order = 132.5, transition bandwidth = 6.25 Hz, cutoff frequency (−6dB) = 28.125 Hz

4 filter order = 1000, transition bandwidth = 0.5 Hz, cutoff frequency (−6dB) = 0.00004 Hz

## Notes

### Competing Interest Statement

The authors have declared no competing interest.

