## Supplementary Material for "The effect of task demand on EEG responses to irrelevant sound and speech in simulated surgical environments"

### 1.1 SURG-TLX items

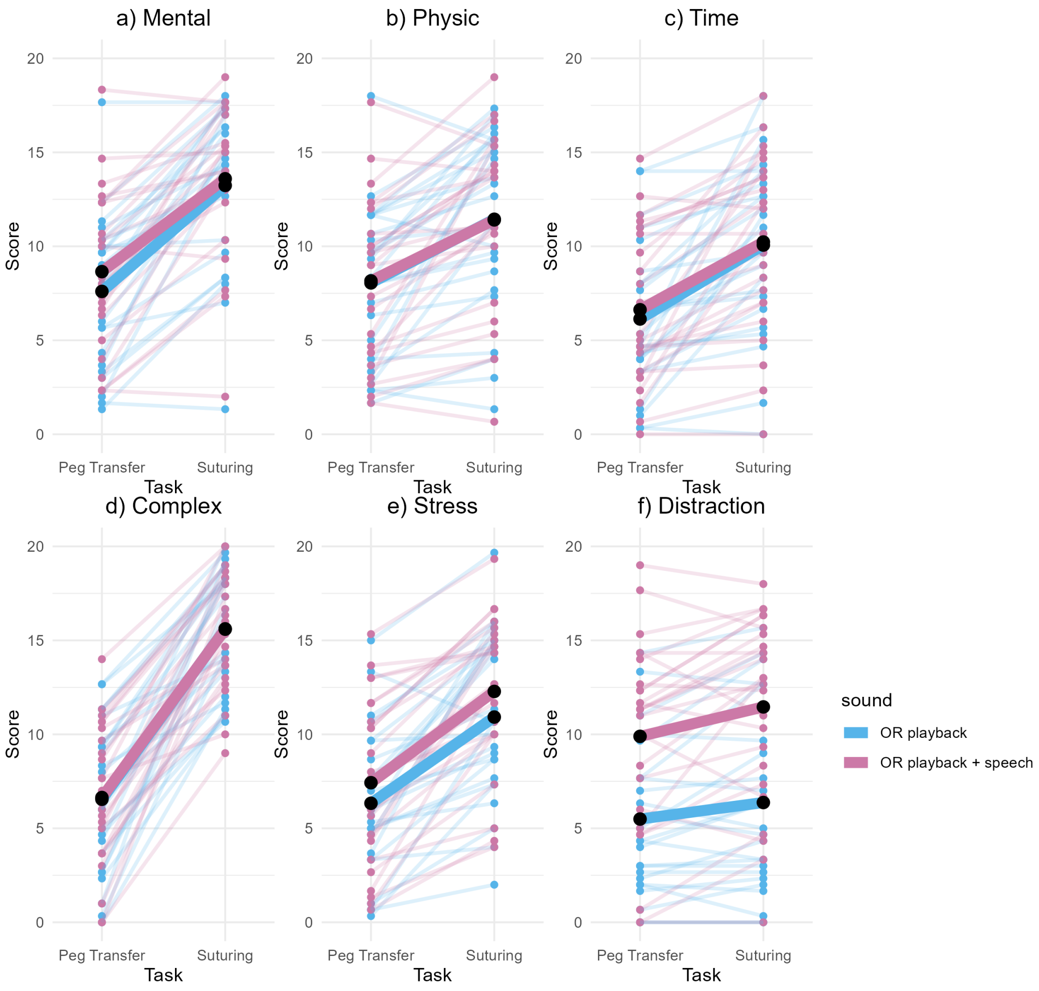


Figure S 1: Score for each item of the SURG-TLX for each task and sound condition. The tasks were selected to represent two difficulty levels, with the peg transfer task representing the easy task and suturing representing the difficult task. Score for each item of the SURG-TLX for each task and sound condition. a)-d) showed a significant effect of task, nut not effect of sound condition or an interaction effect. e)-f) showed a significant effect of task and sound condition, but no interaction effect. The thin lines show the participants' data for each sound condition, the thick line the average across participants.

### 1.2 Model comparisons

Table S 1: *self-reports (except distraction)*

| **Outcome** | **Model Comparison** | **AIC** | **Chisq** | **p-value** | **Fixed Effects of final model (Estimate ± SE)** |
|---|---|---|---|---|---|
| **Model Descriptions:** | | | | | |
| M0: $\hat{y} \sim 1 + (1\|participant)$ | | | | | |
| M1: $\hat{y} \sim task + (1\|participant)$ | | | | | |
| M2: $\hat{y} \sim task + sound(1\|participant)$ | | | | | |
| M3: $\hat{y} \sim task * sound + (1\|participant)$ | | | | | |
| M4: $\hat{y} \sim task + (task\|participant)$ | | | | | |
| M5: $\hat{y} \sim task + sound + (task\|participant)$ | | | | | |
| M6: $\hat{y} \sim task + sound + (task + sound\|participant)$ | | | | | |
| Total | M1 vs M0 | 473.01 | 85.86 | $< 2.2e-16$ *** | Intercept: 6.69 ± 0.70 *** |
| | M2 vs M1 | 462.99 | 12.01 | 0.0005 *** | Task (SU): 4.58 ± 0.50 *** |
| | M3 vs M2 | 464.99 | 0.0001 | 0.9934 | Sound (Sp): 1.20 ± 0.27 *** |
| | M5 vs M2 | 442.31 | 24.68 | 4.37e-06 *** | |
| | **M6** vs M5 | 438.05 | 10.26 | 0.016 * | |
| Mental | M1 vs M0 | 515.32 | 75.04 | $< 2.2e-16$ *** | Intercept: 8.13 ± 0.77 *** |
| | M2 vs M1 | 514.92 | 2.39 | 0.1219 | Task (SU): 5.30 ± 0.66 *** |
| | **M4** vs M1 | 504.79 | 14.52 | 0.0007 *** | |
| Physical | M1 vs M0 | 503.28 | 47.41 | 5.75e-12 *** | Intercept: 8.11 ± 0.82 *** |
| | M2 vs M1 | 505.26 | 0.02 | 0.8965 | Task (SU): 3.33 ± 0.64 *** |
| | **M4** vs M1 | 471.30 | 35.98 | 1.54e-08 *** | |
| Time | M1 vs M0 | 508.78 | 52.86 | 3.58e-13 *** | Intercept: 6.38 ± 0.77 *** |
| | M2 vs M1 | 510.09 | 0.69 | 0.4054 | Task (SU): 3.80 ± 0.68 *** |
| | **M4** vs M1 | 478.05 | 34.73 | 2.87e-08 *** | |
| Complexity | M1 vs M0 | 495.29 | 143.07 | $< 2.2e-16$ *** | Intercept: 6.59 ± 0.75 *** |
| | M2 vs M1 | 497.27 | 0.02 | 0.9025 | Task (SU): 9.03 ± 0.74 *** |
| | **M4** vs M1 | 419.71 | 79.58 | $< 2.2e-16$ *** | |
| Stress | M1 vs M0 | 517.15 | 65.43 | 6.04e-16 *** | Intercept: 6.25 ± 0.79 *** |
| | M2 vs M1 | 511.49 | 7.66 | 0.0056 ** | Task (SU): 4.75 ± 0.62 *** |
| | M3 vs M2 | 513.35 | 0.131 | 0.716 | Sound (Sp): 1.25 ± 0.43 ** |
| | M5 vs M2 | 501.91 | 13.57 | 0.0011 ** | |
| | **M6** vs M5 | 493.67 | 14.25 | 0.0026 ** | |

*Model comparisons for the SURG-TLX total and individual scores. The distraction item can be found in the next table, as the model computation differed for this item from that of the other items. The fixed effects are estimated from the final model, marked in bold. Model descriptions provided at the top of the table. Note that M4 was only tested, if M2 was not significant. SU: Suturing, Sp: Speech present. Significance levels: * $p < .05$, ** $p < .01$, *** $p < .001$.*

Table S 2: *self-report (distraction item)*

| Outcome | Model Comparison | AIC | Chisq | p-value | Fixed Effects of final model (Estimate ± SE) |
|---|---|---|---|---|---|
| **Model Descriptions:** | | | | | |
| M0: $\hat{y} \sim 1 + (1\|participant)$ | | | | | |
| M1: $\hat{y} \sim sound + (1\|participant)$ | | | | | |
| M2: $\hat{y} \sim sound + task + (1\|participant)$ | | | | | |
| M3: $\hat{y} \sim sound * task + (1\|participant)$ | | | | | |
| M4: $\hat{y} \sim sound + task + (sound\|participant)$ | | | | | |
| M5: $\hat{y} \sim sound + task + (sound + task\|participant)$ | | | | | |
| | M1 vs M0 | 534.64 | 60.30 | 8.15e-15 *** | Intercept: 5.30 ± 0.92 *** |
| | M2 vs M1 | 529.86 | 6.78 | 0.0092 ** | Sound (Sp): 4.79 ± 0.48 *** |
| Distraction | M3 vs M2 | 531.20 | 0.66 | 0.4151 | Task (SU): 1.27 ± 0.33 *** |
| | **M4** vs M2 | 514.49 | 19.37 | 6.24e-05 *** | |
| | M5 vs M4 | 518.20 | 2.30 | 0.513 | |

*Model comparisons for the SURG-TLX item distraction. Model descriptions provided at the top of the table. The fixed effects are estimated from the final model, marked in bold. SU: Suturing, Sp: Speech present. Significance levels: * $p < .05$, ** $p < .01$, *** $p < .001$.*

Table S 3: *Surgical task performance*

| Outcome | Model Comparison | AIC | Chisq | p-value | Fixed Effects of final model (Estimate ± SE) |
|---|---|---|---|---|---|
| **Model Descriptions:** | | | | | |
| M0: $\hat{y} \sim 1 + (1\|participant)$ | | | | | |
| M1: $\hat{y} \sim sound + (1\|participant)$ | | | | | |
| PT-Transfers | M1 vs **M0** | 281.80 | 0.0207 | 0.886 | Intercept: 23.013 ± 1.026 *** |
| PT-Drops | M1 vs **M0** | 248.69 | 0.1271 | 0.721 | Intercept: 7.1 ± 0.6 *** |
| SU-Duration | M1 vs **M0** | -166.98 | 0.2195 | 0.64 | Intercept: 2.35 ± 0.26 *** |
| SU-Damage | M1 vs **M0** | 456.49 | 2.2687 | 0.132 | Intercept: 52.673 ± 5.016 *** |

*Model comparisons for the surgical task performance. Model descriptions provided at the top of the table. The fixed effects are estimated from the final model, marked in bold. PT: Peg transfer. SU: Suturing. Significance levels: * $p < .05$, ** $p < .01$, *** $p < .001$.*

Table S 4: *ERP: Presence of gating*

| Outcome | Model Comparison | AIC | Chisq | p-value | Fixed Effects of final model (Estimate ± SE) |
|---|---|---|---|---|---|
| **Model Descriptions:** | | | | | |
| M0: $\hat{y} \sim 1 + (1\|participant)$ | | | | | |
| M1: $\hat{y} \sim position + (1\|participant)$ | | | | | |
| M2: $\hat{y} \sim position + (click\|participant)$ | | | | | |
| amp (Sa) | M1 vs M0 | 248.89 | 29.066 | 6.995e-08 *** | Intercept: 2.97 ± 0.346 *** |
| | **M2** vs M1 | 241.87 | 11.017 | 0.004 ** | position (2nd): -1.16 ± 0.23 *** |
| amp (Sp) | M1 vs M0 | 248.35 | 7.8145 | 0.005 ** | Intercept: 1.74 ± 0.227 *** |
| | M2 vs **M1** | 249.68 | 2.665 | 0.263 | position (2nd): -0.598 ± 0.21 ** |

*Model comparisons to check whether a gating effect is present. The N1-P2 peak-to-peak amplitude was estimated seperately for the 'speech present' and 'speech absent' condition. Model descriptions provided at the top of the table. The fixed effects are estimated from the final model, marked in bold. Sa: Speech absent, Sp: Speech present. Significance levels: * $p < .05$, ** $p < .01$, *** $p < .001$.*

Table S 5: *ERP: Gating difference between tasks*

| Outcome | Model Comparison | AIC | Chisq | p-value | Fixed Effects of final model (Estimate ± SE) |
|---|---|---|---|---|---|
| **Model Descriptions:** | | | | | |
| M0: $\hat{y} \sim 1 + (1\|participant)$ | | | | | |
| M1: $\hat{y} \sim task + (1\|participant)$ | | | | | |
| M2: $\hat{y} \sim task + (task\|participant)$ | | | | | |
| gating (Sa) | **M1** vs M0 | 130.24 | 6.752 | 0.0093 ** | Intercept: 0.764 ± 0.27 ** |
| | M2 vs M1 (did not converge) | | | | Task (SU): 0.79 ± 0.29 * |
| gating (Sp) | M1 vs **M0** | 140.40 | 2.48 | 0.115 | Intercept: 0.6 ± 0.2537 * |

*Model comparisons to investigate whether the strength of gating differed between tasks. Model descriptions provided at the top of the table. The fixed effects are estimated from the final model, marked in bold. PT: Peg transfer, SU: Suturing, Sa: Speech absent, Sp: Speech present. Significance levels: * $p < .05$, ** $p < .01$, *** $p < .001$.*

Table S 6: *TRF: Difference between tasks*

| Outcome | Model Comparison | AIC | Chisq | p-value | Fixed Effects of final model (Estimate ± SE) |
|---|---|---|---|---|---|
| **Model Descriptions:**<br>M0: $\hat{y} \sim 1 + (1\|participant)$<br>M1: $\hat{y} \sim task + (1\|participant)$<br>M2: $\hat{y} \sim task + (task\|participant)$ | | | | | |
| playback response (Sa) | M1 vs **M0** | -183.96 | 2.41 | 0.121 | Intercept: 0.062 ± 0.006 *** |
| playback response (Sp) | M1 vs **M0** | -189.77 | 0.41 | 0.524 | Intercept: 0.045 ± 0.004 *** |
| speech response (Sp) | **M1** vs M0<br>M2 vs M1<br>(did not converge) | -181.39 | 11.72 | 0.001 *** | Intercept: 0.11 ± 0.01 ***<br>Task (SU): -0.02 ± 0.004 *** |

*Model comparisons to investigate whether the tasks influenced the processing of the different continuous stimuli. Model descriptions provided at the top of the table. The fixed effects are estimated from the final model, marked in bold. SU: Suturing, Sa: Speech absent, Sp: Speech present. Significance levels: * $p < .05$, ** $p < .01$, *** $p < .001$.*

Table S 7: ***TRF: prediction of workload***

| Outcome | rc | Model Comparison | AIC | Chisq | p-value | Fixed Effects of final model (Estimate ± SE) |
|---|---|---|---|---|---|---|
| **Model Descriptions:**<br>M0: $\hat{y} \sim task + (1\|participant)$<br>M1: $\hat{y} \sim task + rc + (1\|participant)$<br>M2: $\hat{y} \sim task * rc + (1\|participant)$ | | | | | | |
| TLX total | Playback response (Sa) | M1 vs **M0**<br>M2 vs M1 | 212.49<br>213.96 | 0.803<br>1.337 | 0.370<br>0.513 | Intercept: 7.02 ± 0.81 ***<br>Task (SU): 4.8 ± 0.68 *** |
| TLX total | Playback response (Sp) | M1 vs **M0**<br>M2 vs M1 | 210.20<br>212.20 | 0.738<br>0.742 | 0.390<br>0.690 | Intercept: 8.35 ± 0.83 ***<br>Task (SU): 4.63 ± 0.61 *** |
| TLX total | speech response (Sp) | M1 vs M0<br>M2 vs **M1** | 206.25<br>208.21 | 4.687<br>0.041 | 0.030 *<br>0.840 | Intercept: 8.67 ± 0.78 ***<br>Task (SU): 3.98 ± 0.63 ***<br>rc: -1.389 ± 0.637 * |

*Model comparisons to investigate whether the centered correlation (rc) between the actual and reconstructed stimulus envelope predict self-reported workload. Model descriptions provided at the top of the table. The fixed effects are estimated from the final model, marked in bold. SU: Suturing, Sa: Speech absent, Sp: Speech present. Significance levels: * $p < .05$, ** $p < .01$, *** $p < .001$.*
